# Aryl Hydrocarbon Receptor Suppresses Eosinophilic Esophagitis Responses through OVOL1 and SPINK7

**DOI:** 10.1101/2023.05.17.541192

**Authors:** Nurit P. Azouz, Andrea M. Klingler, Mark Rochman, Misu Paul, Julie M. Caldwell, Michael Brusilovsky, Alexander T. Dwyer, Xiaoting Chen, Daniel Miller, Arthur Lynch, Carmy Forney, Leah C. Kottyan, Matthew T. Weirauch, Marc E. Rothenberg

**Affiliations:** Division of Allergy and Immunology, Cincinnati Children’s Hospital Medical Center, Department of Pediatrics, University of Cincinnati College of Medicine, Cincinnati, Ohio, USA; Department of Pediatrics, University of Cincinnati College of Medicine, Cincinnati, OH, USA; Center for Stem Cell and Organoid Medicine, Cincinnati Children’s Hospital Medical Center, Cincinnati, Ohio, USA; Division of Pediatric General and Thoracic Surgery, Cincinnati Children’s Hospital Medical Center, Cincinnati, Ohio, USA; Center for Autoimmune Genomics and Etiology, Cincinnati Children’s Hospital Medical Center, University of Cincinnati College of Medicine, Cincinnati, Ohio, USA; Divisions of Human Genetics, Biomedical Informatics, and Developmental Biology, Cincinnati Children’s Hospital Medical Center, University of Cincinnati College of Medicine, Cincinnati, Ohio, USA

**Keywords:** eosinophilic esophagitis (EoE), aryl hydrocarbon receptor (AHR), ovo like transcriptional repressor 1 (OVOL1), interleukin-13 (IL-13), calpain-14

## Abstract

Eosinophilic esophagitis (EoE) is a type 2 allergic disease characterized by esophageal inflammation and epithelial cell dysfunction. Acquired loss of the anti-serine protease of kazal type 7 (SPINK7) in the squamous epithelium of the esophagus has a causal role in EoE pathogenesis. Yet there is a limited understanding of the factors that regulate its expression and responsiveness to inflammatory stimuli. Herein, we identified the transcription factor, ovo like transcriptional repressor 1 (OVOL1) as an esophageal selective gene product that regulates SPINK7 promoter activity. Overexpression of *OVOL1* increased *SPINK7* expression, whereas, its depletion decreased *SPINK7* expression, impaired epithelial barrier and increased production of the pro-atopy cytokine thymic stromal lymphopoietin (TSLP). Mechanistically, ligands of AHR induced nuclear translocation of OVOL1 which in turn promoted epithelial cell differentiation, barrier function and *SPINK7* expression. Interleukin (IL)-4 and IL-13 abolished AHR ligand-induced OVOL1 nuclear translocation. Stimulation with IL-13 abrogated the nuclear translocation of OVOL1 and promoted enhanced degradation of OVOL1 protein. This effect of IL-13 was dependent on the esophageal specific cysteine protease calpain-14. Translational studies demonstrated loss of OVOL1 protein expression in patients with EoE. In summary, AHR mediates its action via OVOL1-induced SPINK7 transcription, and IL-4 and IL-13 repress this pathway in EoE. As such, activation of the AHR pathway is a potential intervention strategy for reversing EoE.

**Graphical abstract:** 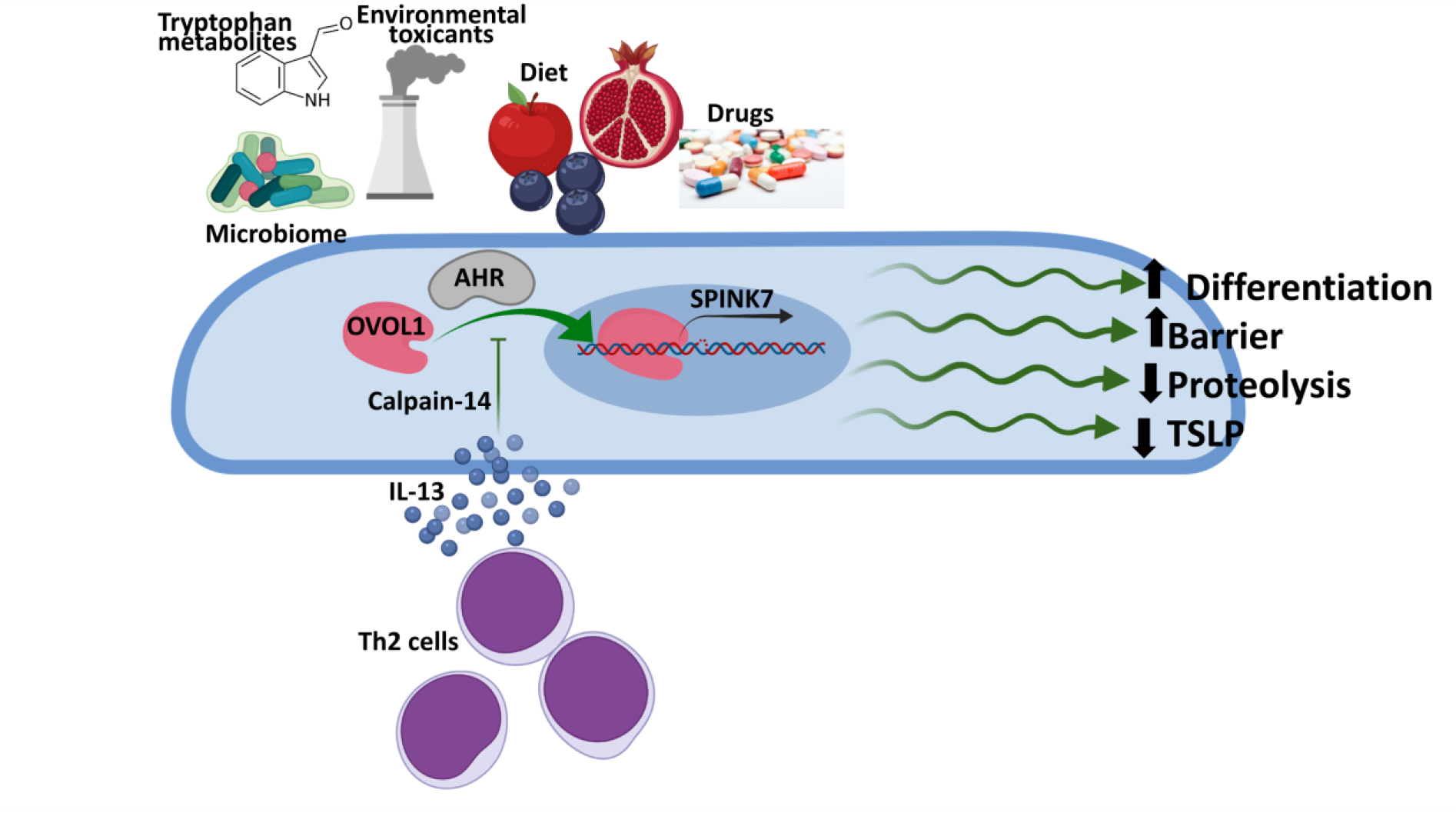

**The influence of the exposome on regulatory networks in EoE pathogenesis.**

AHR is activated and influenced by diet nutrients, environmental toxicants, microbiome composition, tryptophan metabolites, and drugs. When AHR is activated, it promotes translocation of OVOL1 to the nucleus, which in turn promotes expression of epithelial genes including *SPINK7*. SPINK7 expression promotes epithelial differentiation, barrier function, decreased proteolytic activity, and decreased TSLP production. IL-4 and IL-13 inhibit OVOL1 nuclear translocation and therefore, repress *SPINK7* expression. IL-13–stimulated *CAPN14* expression decreases OVOL1 protein expression and *SPINK7* transcription.

## Introduction

EoE is a chronic, food antigen–driven disease of the esophagus (1). Although EoE is considered an orphan disease, 160,000 people are diagnosed with EoE in the United States. The increased prevalence of EoE and other allergic diseases have been attributed to changes in environmental factors, particularly those associated with dysbiosis of commensal flora (3–12). Mechanistically, EoE is mediated by interleukin (IL) 13 overexpression, which subsequently induces CCL26 production that recruits eosinophils and overproduction of the cysteine protease calpain 14 (13–16).

In the squamous epithelium of the skin and esophagus, expression of the anti-serine proteases of the kazal type (SPINK) maintains homeostatic control of inflammation. Loss of SPINK5 and/or SPINK7 (the two main SPINK family members expressed in the squamous epithelium), leads to profound consequences including impaired epithelial barrier function and elicitation of allergic inflammation in the skin and/or esophagus (17–20). Depletion of SPINK7 in esophageal epithelial cells is sufficient to induce barrier dysfunction, elicits marked production of pro-inflammatory and pro-atopy cytokines including that the alarmin thymic stromal lymphopoietin (TSLP), that is encoded by the gene involved in a primary EoE susceptibility locus (21), and activation of eosinophils by the urokinase plasminogen activator pathway (17). Rare homozygous mutations of *SPINK5* as well as *Spink5* deletion in mice, are sufficient to elicit loss of epithelial barrier integrity, and pro-inflammatory and pro-atopy responses including atopic dermatitis and EoE in vivo (22–24). Yet, there is little data about how these SPINK proteins are regulated under basal conditions, and how environmental sensing can lead to their loss.

Herein, we interrogated the regulatory mechanisms that control *SPINK7* expression during epithelial differentiation. We identified elevated histone 3 acetylation marks at the *SPINK7* promoter region in the cells during differentiation. Binding sites for the C2H2 zinc finger transcription factor, ovo like transcriptional repressor 1 (OVOL1) are found in the SPINK7 promoter region and are co-localized with the histone 3 acetylation marks. Binding of OVOL1 to the SPINK7 promoter was required for its activity. Mechanistically, a variety of AHR ligands (e.g., proton-pump inhibitors, dietary compounds, metabolites produced by bacteria and particulate matter), induced OVOL1 nuclear localization and subsequent *SPINK7* expression. Furthermore, the type 2 cytokines IL-4 and IL-13, which are inducers of allergic responses, repressed OVOL1 activation. Additionally, the product of the chief EoE susceptibility locus (2p23) calpain-14 (25), an intracellular regulatory protease induced by IL-13 in esophageal epithelial cells (16), resulted in post-transcriptional modification of OVOL1 levels. Translational studies demonstrated a marked loss of OVOL1 protein expression in esophageal biopsies of EoE patients compared to control patients. We suggest that AHR serves as an esophageal sensor for environmental signals and has the potential to rapidly control esophageal epithelium fate by altering SPINK7 levels via OVOL1 activation. As such, modulation of this pathway (e.g., supplementing AHR ligands) may be therapeutic in EoE and related allergic diseases.

## Results

### SPINK7 expression is induced during epithelial differentiation

We analyzed *SPINK7* expression in cultures of esophageal epithelial progenitor cells (EPC2 cells) under conditions that promote cellular differentiation, namely high confluency and high calcium. *SPINK7* expression was more strongly induced under high-confluency conditions compared to low-confluency conditions (Figure 1A). *SPINK7* expression was further increased under high-calcium (1.8 mM) conditions compared to low-calcium conditions (0.09 mM; Figure 1A). Epigenetic analysis of the *SPINK7* promoter region of EPC2 cells revealed an increase in H3K27 acetylation marks in the region surrounding *SPINK7* transcription start site (TSS) in high-confluency conditions compared to low-confluency conditions (Figure 1B). The highest levels for the K27 acetylation mark were observed in cells that were grown in high confluency and high calcium (1.8 mM; Figure 1B).

**Figure 1.**
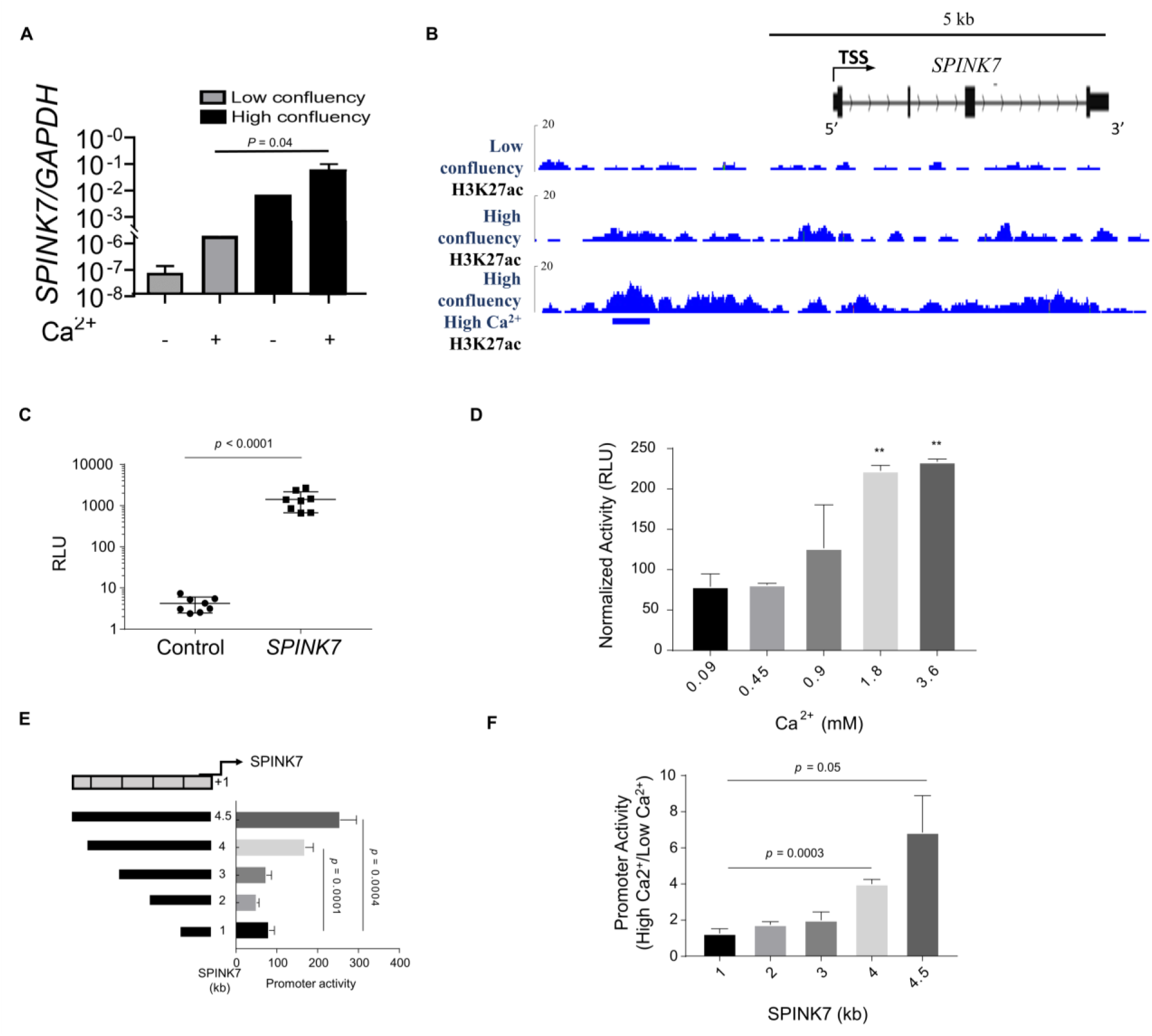
*SPINK7* expression is induced by calcium and cell confluency. **A.** Quantitative polymerase chain reaction (qPCR) of *SPINK7* expression in EPC2 in cells. **B.** Levels of H3K27Ac in the promoter region of *SPINK7* in EPC2 cells in the indicated conditions. **C.** Promoter activity in cells grown in high calcium and high confluency, co-transfected with either nano-luciferase (nLUC) vector containing the *SPINK7* promoter (*SPINK7*) or a promoterless nLUC vector and with firefly vector to control for transfection efficiency (Control), presented as relative luminescence units (RLU). **D.** Promoter activity in cells co-transfected with SPINK7-nLUC that were grown in the indicated concentrations of CaCl_2_ and normalized to cells co-transfected with nLUC and firefly vector. **E.** Promoter activity in cells that were grown in 1.8 mM of CaCl_2_ and co-transfected with nLUC constructs that contain either 0, 1, 2, 3, 4, or 4.5 kb of the *SPINK7* promoter sequence and firefly vector. **F.** Promoter activity in cells co-transfected with nLUC constructs, containing either 0, 1, 2, 3, 4, or 4.5 kb of the *SPINK7* promoter sequence and firefly vector, and grown in either 0.09 or 1.8 mM of CaCl_2_. The values of the 1.8 mM of CaCl_2_ lysates were divided to the values of the cells cultured in 0.09 mM of CaCl_2_. A and C-F, data represent mean ± SEM.

### Identification of the *SPINK7* promoter region

We identified a regulatory region in the 4.5-kb 5’ region flanking the *SPINK7* TSS using transcriptional and epigenetic data (Figure 1B). To test the activity of this putative promoter, EPC2 cells were grown at high density and in high calcium (180 µM) conditions to induce cell differentiation. Then, cells were transiently co-transfected with firefly luciferase vector (to control for transfection efficiency) and either a reporter construct containing the presumed promoter region (the 4.5-kb 5’ region flanking the *SPINK7* TSS, referred to henceforth as *SPINK7*) or a control promoterless vector. The *SPINK7*-transfected cells had on average an approximately 340-fold increase in luminescence signal compared to cells transfected with the control promoterless vector (p < 0.0001; Figure 1C). Calcium ions increased *SPINK7* promoter activity in a dose-dependent manner and reached a plateau at 1.8 mM of calcium (Figure 1D). No difference was observed in the control promoterless vector activity in different calcium concentrations (Supplementary Figure 1A). These collective data indicate that this regulatory region of the *SPINK7* gene has calcium-dependent promoter activity.

We subsequently determined the minimal sequence required for promoter activity. The promoter activity of all reporter constructs was sufficient to drive at least some activity (Figure 1E). The luciferase activity of the cells transfected with the 4.5-kb and 4-kb constructs increased by 3-fold and 2-fold, respectively, compared to the luciferase activity of the cells that were transfected with the 1-kb construct, (p = 0.0004, p = 0.0001 respectively; Figure 1E). The luciferase activity of the cells transfected with the 3-kb and 2-kb constructs was not significantly different than the luciferase activity of the cells transfected with the 1-kb construct (Figure 1E). In low-calcium conditions, low promoter activity was observed in all of the constructs, with no difference between the 1-kb and the 4.5-kb or 4-kb constructs (Supplementary Figure 1B). For the 4.5-kb and 4-kb constructs, luciferase activities were increased by 7-fold and 4-fold, respectively, in the high-calcium compared to the low-calcium media (p = 0.05 and p = 0.0003, respectively; Figure 1F). The 3-kb, 2-kb, and 1-kb constructs were not affected by high calcium (Figure 1F). These collective data suggest that induction of cellular differentiation (i.e., high-calcium and high-confluency conditions) promotes *SPINK7* promoter activity via the 4.5-kb region upstream of the *SPINK7* TSS.

### The transcription factor OVOL1 regulates *SPINK7* expression

We next asked which transcription factors (TFs) regulate SPINK7 promoter activity. Analysis of the *SPINK7* promoter using TF binding motifs obtained from the CisBP database (build 1.02) (26) revealed several TFs that are predicted to bind the *SPINK7* promoter. We narrowed this list of TFs to 36 by intersecting with genes that are induced during esophageal epithelial differentiation (as previously reported by us (17)) or with esophageal specific genes or with genes that are dysregulated in patients with EoE compared to control patients (Supplementary Table 1). We transiently overexpressed 4 resulting candidate TFs that may regulate *SPINK7* expression (i.e., OVOL1, VDR, POU2F3, and PRDM1). Overexpression of VDR (in the presence or absence of the VDR ligand calcipotriol), POU2F3, and PRDM1 did not increase *SPINK7* promoter activity (Figure 2A). In contrast, overexpression of OVOL1 increased *SPINK7* promoter activity by 2.4 fold (p = 0.0006; Figure 2A). Western blot analysis confirmed that OVOL1 protein was overexpressed in the *OVOL1*-transfected cells compared to control cells (Supplementary Figure 2A).

**Figure 2.**
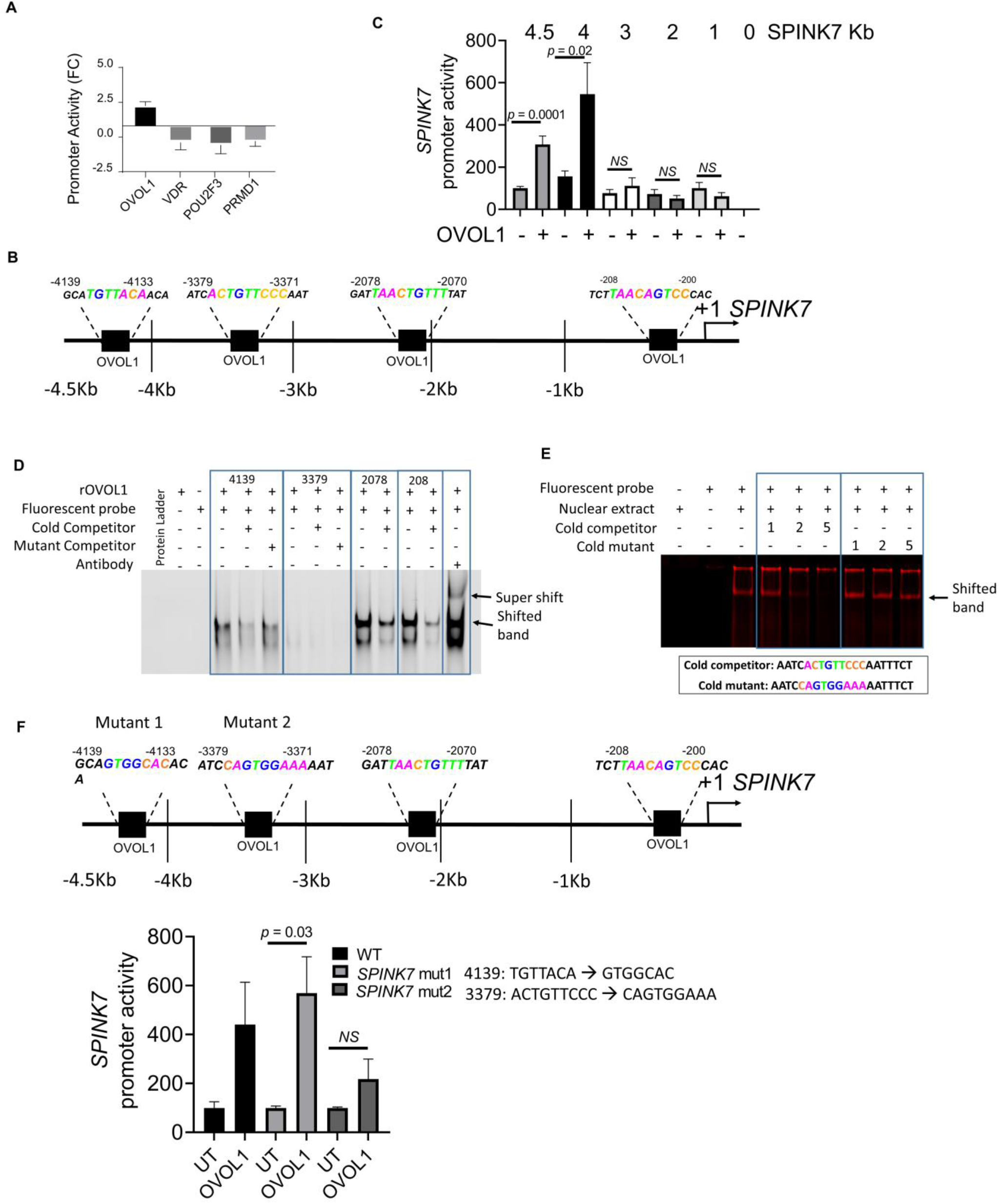
OVOL1 binds to the SPINK7 promoter and induces *SPINK7* expression. **A.** Nano-luciferase activity in lysates from cells transfected with *SPINK7* promoter, firefly plasmid, and plasmids encoding for the indicated transcription factors normalized to control lysates from cells transfected with *SPINK7* promoter, firefly plasmid, and an empty plasmid. **B.** Analysis of the proximal human *SPINK7* promoter sequence identified four potential OVOL1 binding sites. Transcription Start Site +1. **C.** Nano-luciferase activity in lysates co-transfected with either OVOL1 or a control plasmid and with *SPINK7* promoter deletion constructs. **D.** Representative results from EMSA experiments using recombinant human OVOL1 protein. Fluorescent IRDye 700–labeled probe sequence 5’-AATCACTGTTCCCAATTTCT-3’; unlabeled (cold) WT competitor sequence: 5’-AATCACTGTTCCCAATTTCT-3’; cold mutant sequence: 5’-AATCCAGTGGAAAAATTTCT-3’; 4139: 5’-CATTTCTGTTACATTAGGAT-3’; 4139 mutant: 5’-CATTTCGTGGCACTTAGGAT-3’; 2078: 5’-GTAGATTAACTGTTTATGTT-3’; 208: 5’-ATTCTTAACAGTCCCACCTT-3’. **E.** Representative results from EMSA experiment using nuclear extracts from HEK-293T cells transfected with OVOL1 plasmid or a control plasmid. Fluorescent IRDye 700–labeled probe sequence 5’-AATCACTGTTCCCAATTTCT-3’; cold competitor sequence: 5’-AATCACTGTTCCCAATTTCT-3’; cold mutant sequence: 5’-AATCCAGTGGAAAAATTTCT-3’. Cold competitors were added in concentrations of 1X, 2X and 5X compared to the fluorescent probe (presented as 1, 2, and 5). **F.** Nano luciferase activity in lysates co-transfected with OVOL1 or a control plasmid and with the *SPINK7* promoter with either mutated OVOL1 binding site 1 (-4,139 bp: 5’-GCATGTTACAACA-3 → 5-GCAGTGGCACACA-3’) or mutated OVOL1 binding site 2 (-3379 bp: 5’-ATCACTGTTCCCAAT-3’ → 5’-ATCCAGTGGAAAAAT-3’) or wild-type *SPINK7* promoter. Cells were either left untreated or treated with FICZ (1 µM). A, C, and F, data represent mean ± SEM. FC, fold change; NS, not significant; UT, untreated; WT, wild type.

### OVOL1 binds to the *SPINK7* promoter and activates *SPINK7* expression

We next examined whether *SPINK7* is a di2rect OVOL1 target gene in esophageal epithelial cells. We predicted 4 OVOL1 binding sites upstream of the *SPINK7* TSS (Figure 2B). We tested the requirement of this response by examining *SPINK7* promoter activity in the presence or absence of *OVOL1* overexpression (Figure 2B). Overexpression of *OVOL1* induced promoter activity in cells that were transfected with the 4-kb and 4.5-kb constructs, with similar promoter activity (Figure 2C). In contrast, the promoter activity of cells that were transfected with the 3-kb, 2-kb or 1-kb *SPINK7* constructs was not significantly affected by *OVOL1* overexpression (Figure 2C).

We analyzed the binding of recombinant human OVOL1 protein to a fluorescent DNA probe corresponding to the -4139, -3379, -2078, and -208 regions of the *SPINK7* promoter by Electrophoretic Mobility-Shift Assay (EMSA). OVOL1 shifted the mobility of the fluorescent probe in all probes, except the -3379 probe (Figure 2D). Unlabelled (cold) wild-type competitors that contain the predicted binding site at -4139, -2078, and -208 bp inhibited the mobility shift (Figure 2D). A mutant cold competitor that contains the predicted binding site at -4139 bp (GTGGCAC) failed to inhibit the mobility shift (Figure 2D). In addition, administration of a rabbit anti-human OVOL1 antibody resulted in a supershift of the -2078 probe. To examine the possibility that additional cellular co-factors are required for the binding of OVOL1, we generated nuclear extracts of HEK-293T cells overexpressing OVOL1. The nuclear extracts shifted the -3379 probe, whereas the cold wild-type probe inhibited the shifted band in a dose-dependent manner (Figure 2E). A mutant cold competitor that contains the predicted binding site at -3379 bp (CAGTGGAAA) failed to inhibit the mobility shift (Figure 2E).

We tested the specificity of this response in two predicted OVOL1 binding sites that affected the *SPINK7*-nLUC activity when absent from the promoter region (Figure 2C). We constructed mutants in these predicted OVOL1 binding sites and assessed *SPINK7*-nLUC promoter reporter construct activation (Figure 2F). *SPINK7*-nLUC activation was significantly decreased for the OVOL1 binding site mutant at -3379 bp when the ACTGTTCCC sequence was replaced with CAGTGGAAA (Figure 2F). In contrast, there was not a significant effect on SPINK7-nLUC activation for the OVOL1 binding site mutant at -4139 bp when the TGTTACA sequence was replaced with GTGGCAC (Figure 2F). These collective data suggest that OVOL1 binds to all four sites in the *SPINK7* promoter and that the -3379 bp region of the *SPINK7* TSS is important for OVOL1-dependent gene expression.

### Loss of *OVOL1* promotes impaired barrier function and TSLP production

We examined the consequences of depleting *OVOL1* expression in EPC2 cells by stable transduction with a vector expressing either shRNA targeting *OVOL1* or non-silencing control (NSC) shRNA (Figure 3A and Supplementary Figure 2B). *OVOL1-*silenced cells that were differentiated in air-liquid interface (ALI) culture system had a 3-fold decrease in *SPINK7* expression compared to that of differentiated NSC-treated cells (Figure 3B). *OVOL1* silenced cells had increased TSLP release compared to NSC-treated cells after polyI:C stimulation (Figure 3C). This is consistent with the phenotype of *SPINK7* depleted cells, which were previously demonstrated to produce increased levels of TSLP upon polyI:C stimulation (17). *OVOL1* silenced cells displayed barrier impairment as assessed by decreased trans epithelial electrical resistance (TEER; Figure 3D). These data suggest that *OVOL1* expression is critical for maintaining *SPINK7* expression, barrier integrity and controlling TSLP production by epithelial cells.

**Figure 3.**
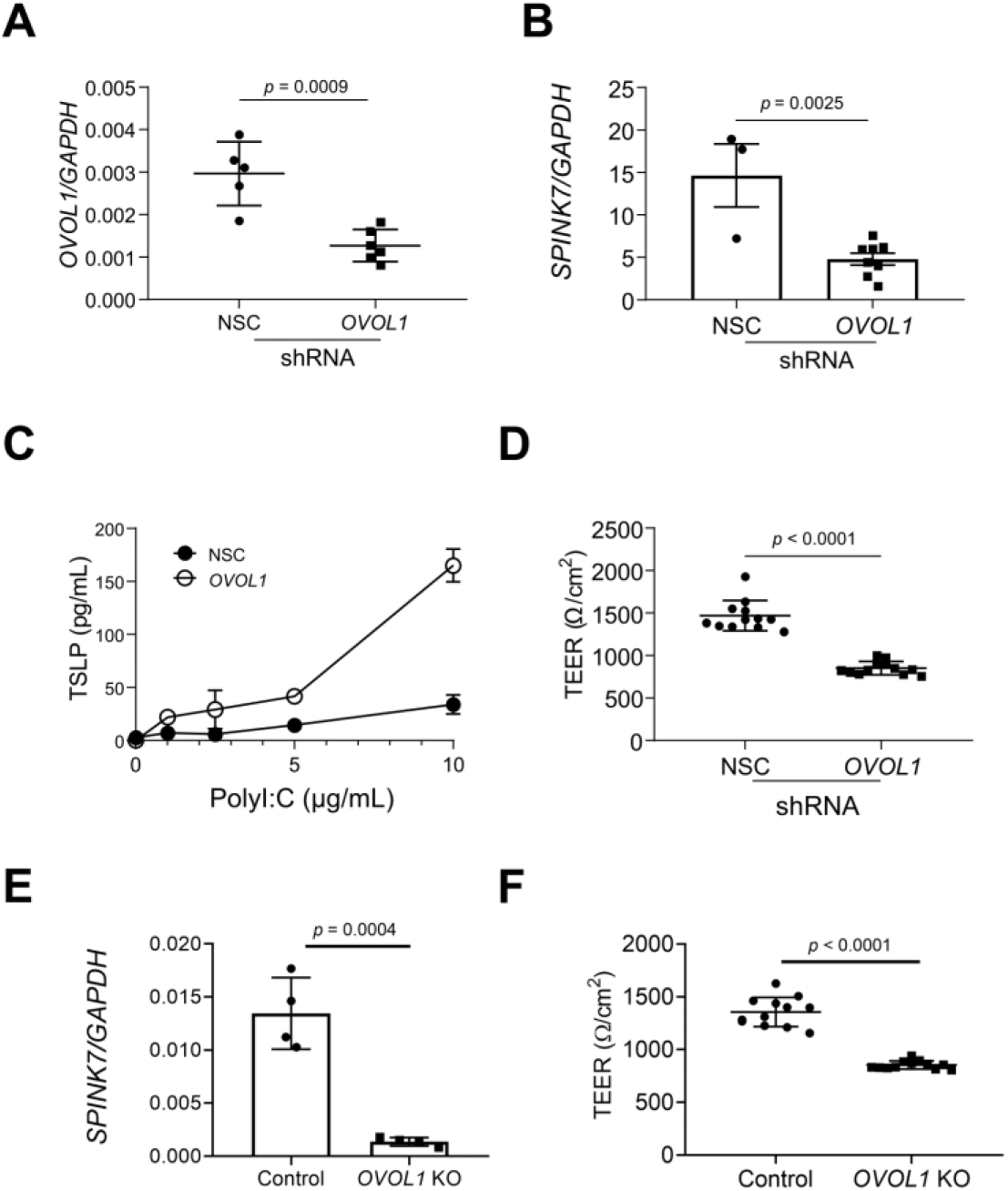
Loss of OVOL1 impairs barrier function and promotes TSLP production. **A.** qPCR analysis of *OVOL1* expression from NSC-treated and *OVOL1*-silenced EPC2 cells. **B.** qPCR analysis of *SPINK7* expression from NSC-treated and *OVOL1*-silenced EPC2 cells at day 14 of ALI differentiation. **C.** TSLP release from NSC-treated and *OVOL1*-silenced EPC2 cells that were grown in high-calcium media for 64 hours and then stimulated for 8 hours with the indicated concentrations of polyinosinic-polycytidylic acid (polyI:C). Cell supernatants were assessed for TSLP levels from three independent experiments. Data are the means ± SD. **D.** TEER (ohm/cm2) measurement from NSC-treated, *OVOL1*-silenced EPC2 cells at day 7 of ALI differentiation. Data are the means ± SD from three independent experiments performed in triplicate. **E.** qPCR analysis of *SPINK7* expression from CRISPR/Cas9 *OVOL1* knockout (KO) and control EPC2 cells at day 14 of ALI differentiation. **F.** TEER (ohm/cm2) measurement from CRISPR/Cas9 *OVOL1* KO and control EPC2 cells at day 7 of ALI differentiation. Data are the means ± SD from three independent experiments performed in triplicate. All P values were calculated by t test (unpaired, two-tailed).

We subsequently generated *OVOL1* knockout cells using clustered regularly interspaced short palindromic repeats (CRISPR)/Cas9 genomic editing (Supplementary Figure 3A-B). *OVOL1* knockout cells that were differentiated in ALI culture had a 10-fold decrease in *SPINK7* expression compared to that of differentiated control EPC2 cells (p = 0.0004; Figure 3E) and had barrier impairment as assessed by decreased TEER (Figure 3F).

### OVOL1 protein expression is lost in EoE

Analysis of *OVOL1* mRNA expression did not reveal any significant difference between EoE biopsies and controls (Figure 4A and Supplementary Table 1). However, we noted that OVOL1 intracellular localization changed by epithelial compartment. In the differentiating cells closer to the basal membrane, OVOL1 localization was nuclear, whereas in the differentiated epithelium (the layers of epithelium near the lumen), OVOL1 localization was cytoplasmic (Figure 4B). Analysis of OVOL1 protein expression revealed a decrease in protein expression in EoE compared to control biopsies (Figure 4C). Western blot analysis of OVOL1 showed that OVOL1 protein expression is decreased by 10-fold in esophageal biopsies from patients with EoE compared to control patients (Figure 4D). Notably, a higher molecular band of OVOL1 (50 kDa) remained evident in control and EoE biopsies (Supplementary Figure 4). These collective data suggest that the OVOL1 protein is deficient in the esophagus of patients with EoE compared to control individuals.

**Figure 4.**
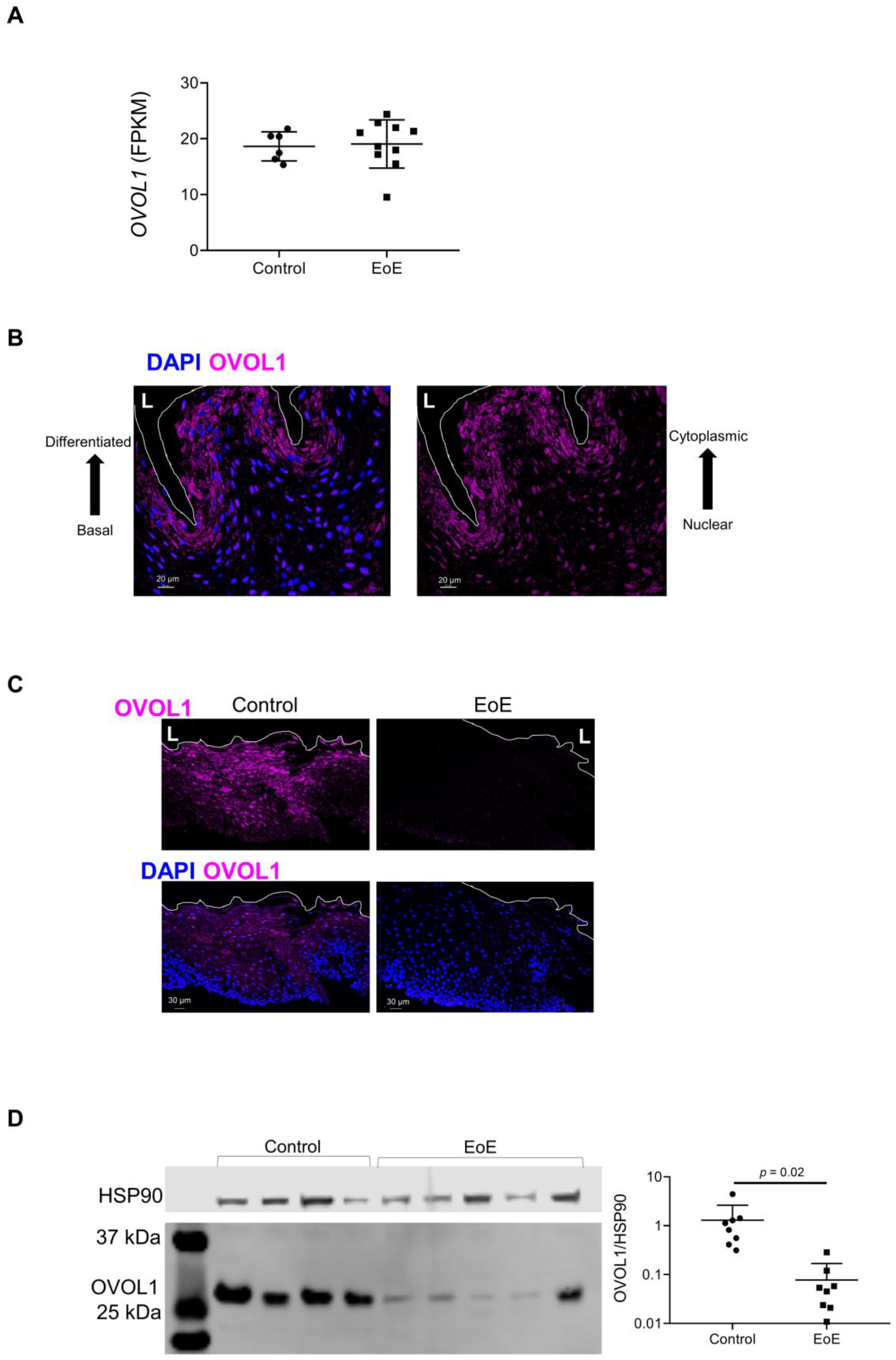
Loss of OVOL1 in EoE biopsies. **A.** mRNA expression of OVOL1 in EoE biopsies compared with control biopsies. **B.** A representative image of immunofluorescence staining of OVOL1 (pink) and DAPI (blue) staining in control biopsy. White line separates the lumen from the epithelium, the lumen side is marked by the letter “L”. **C.** Representative images of immunofluorescence staining of OVOL1 (pink) and DAPI (blue) staining in control and EoE biopsies. White line separates the lumen from the epithelium and the lumen side is marked by the letter “L”. **D.** Western blot analysis of OVOL1 expression in control and EoE biopsies. The graph on the right shows the OVOL1 expression relative to HSP90.

### Environmental and internal cues promote *SPINK7* expression in vitro

Activation of AHR has been demonstrated to promote nuclear localization and activation of OVOL1 in skin keratinocytes (27, 28). AHR is activated in response to a variety of ligands, such as environmental toxicants, including vehicle exhaust and cigarette smoke; dietary compounds (i.e., flavonoids, indole-3-carbinol derivatives, extracts from fruits, vegetables especially cruciferous); products from commensal bacteria (e.g., 6-Formylindolo[3,2-b]carbazole [FICZ], kynurenine and Butyrate); tryptophan metabolism; and drugs including, proton pump inhibitors, which are interestingly used to treat EoE. Different ligands can promote different signaling pathways through AHR (29, 30). Analysis of *AHR* expression in EoE compared to control biopsies revealed that *AHR* is significantly increased in EoE (Supplementary Figure 5). Interestingly, the AHR-regulated genes *NQO1*, *HMOX1*, and *HMOX2* were found to be significantly downregulated in EoE compared to control biopsies (Supplementary Figure 5), demonstrating that AHR signaling is likely dysregulated in EoE.

We subsequently tested the hypothesis that activators of the AHR pathway might regulate *SPINK7* expression via OVOL1. We stimulated EPC2 cells with the AHR ligands FICZ, Omeprazole, urolithin A (UroA), 2-(1’ H-indole-3’-carbonyl)-thiazole-4-carboxylic acid methyl ester (ITE), quercitin, and benzo[a]pyrene (B[a]P). FICZ, Omeprazole, UroA, ITE, and B[a]P efficiently stimulated the expression of the AHR target gene *CYP1A1* and *SPINK7* expression (Figure 5A-B). We next aimed to determine whether *SPINK7* upregulation was AHR dependent. Accordingly, cells pre-treated with the AHR antagonist N-(2-(3H-indol-3-yl)ethyl)-9-isopropyl-2-(5-methyl-3-pyridyl)-7H-purin-6-amine (GNF-351) demonstrated significantly decreased *SPINK7* upregulation induced by all ligands that induced *SPINK7* expression except ITE (Figure 5B). Next, we investigated whether the AHR ligands promote OVOL1 nuclear translocation as previously demonstrated in skin keratinocytes (27). FICZ and ITE induced OVOL1 nuclear mobilization after 1 h of stimulation, whereas 1 h of Omeprazole stimulation was not sufficient to promote OVOL1 nuclear mobilization as assessed by immunostaining (Figure 5C). By 18 hours, Omeprazole induced nuclear mobilization of OVOL1 (Figure 5C). Nuclear fractionation of the cells revealed that the nuclear portion of OVOL1 was increased following FICZ stimulation (Figure 5D).

**Figure 5.**
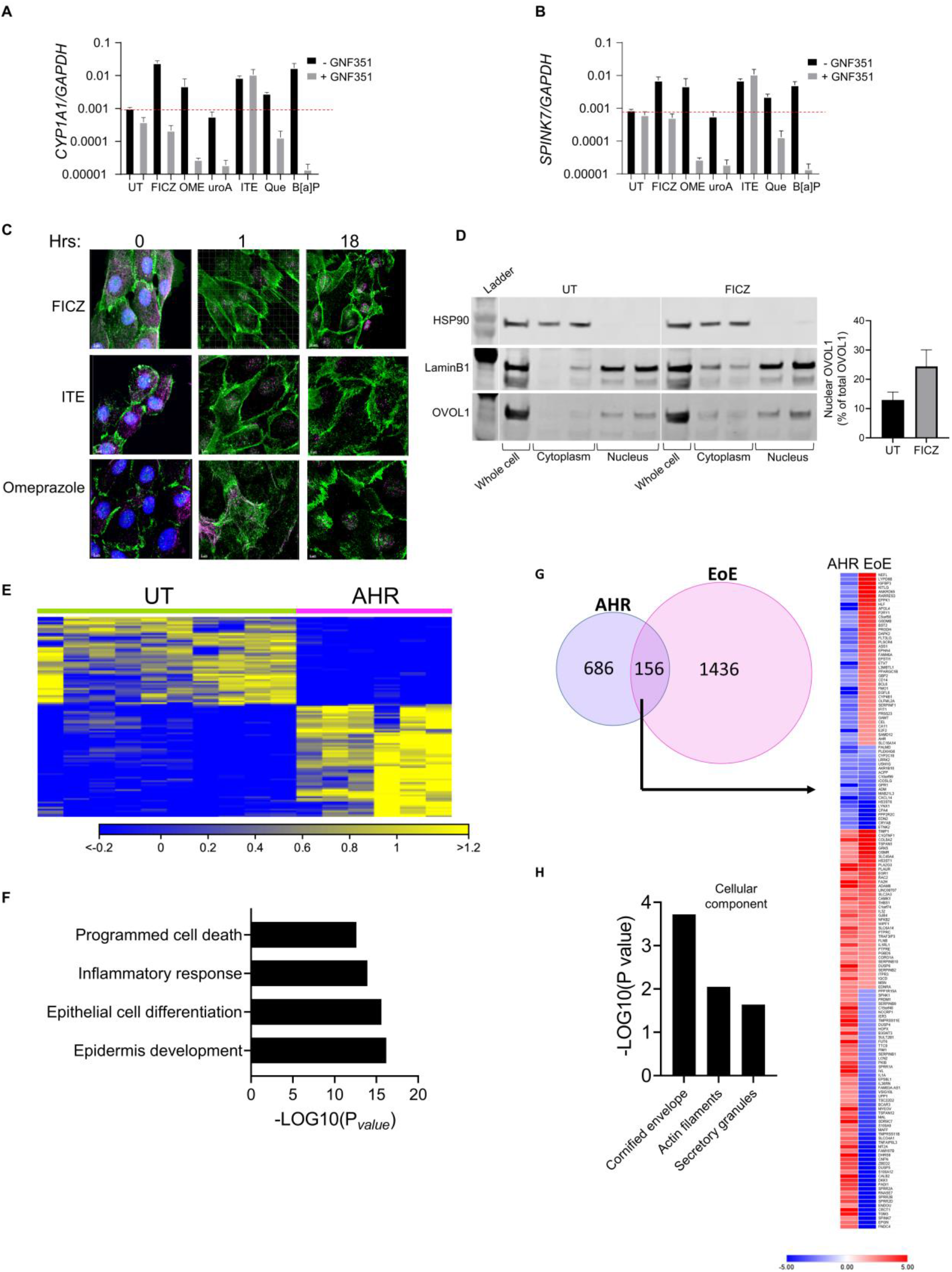
Environmental cues affect *SPINK7* expression in an AHR-dependent and AHR-independent manner. qPCR analysis of *CYP1A1* **(A)** and *SPINK7* **(B)** expression after stimulation with the indicated stimuli with or without GNF351. A and B, red dashed line represents the basal expression of *CYP1A1* and *SPINK7* in UT cells respectively. **C.** Representative images of co-immunofluorescence of desmoglein1 (DSG1, green), OVOL1 (pink) and DAPI (blue) after 0, 1 or 18 hrs of stimulation with FICZ (1 µm), ITE (1 µm), or Omeprazole (10 µm). **D.** Representative western blot of OVOL1 expression in cell fractions. LaminB1 represents a nuclear loading control and HSP90 represents a cytosolic loading control. **E.** Heatmap representing TPM values of genes that are significantly altered by FICZ treatment (p adjusted < 0.05). **F.** Gene ontology (GO) enrichment analyses of genes that are induced by FICZ treatment. P value for GO analysis was calculated by an ANOVA test. **G.** Genes differentially expressed (fold change > I2I) in EPC2 cells following 18 hrs of FICZ stimulation (AHR transcriptome) compared with untreated cells (left column; AHR) compared to the EoE transcriptome (right column; EoE) obtained from RNA sequencing of esophageal biopsies (n = 6 control patients [Control] and n = 10 patients with active EoE [EoE] as described in (59)). Venn diagrams compare the genes of the AHR transcriptome and the EoE transcriptome. Heatmap represents the fold change of genes that overlapped between the FICZ transcriptome and the EoE transcriptome. **H.** GO analysis depicting cellular components of the overlapping genes. The p value for GO analysis was calculated by ANOVA. UT, untreated.

### Consequences of AHR stimulation of esophageal epithelium

Genome-wide transcriptomic analysis of cells stimulated with the AHR ligand FICZ revealed 842 dysregulated genes (i.e., AHR transcriptome; Figure 5E, Supplementary Table 2 and supplementary information). This list of genes included enzymes that are established AHR targets such as *CYP1A1*, NAD(P)H Quinone Dehydrogenase 1 *(NQO1)* and Heme Oxygenase 2 *(HMOX2)*. The up-regulated genes (421 genes) were enriched in functional pathways involved in epidermis development and epithelial differentiation, such as *IVL* (encoding for the barrier gene and differentiation marker involucrin); *TGM3* and *TGM5* (Transglutaminases that catalyze the crosslinking of proteins in the cornified envelop); late cornified envelope protein family including, *LCE3E* and *LCE3D;* and the keratins *KRT9, KRT34 and KRT80* (Figure 5F and Supplementary Table 2). FICZ-upregulated genes were also enriched for regulators of inflammatory responses including the phosphatase *PTPRC* and protease inhibitors (i.e., *SPINK7*, *SERPINB1*, *SERPINB9*, *TIMP1* and *TIMP3*; Figure 5F). A substantial subset (18%) of the FICZ-dysregulated genes overlapped with the EoE transcriptome (156 genes out of 842 FICZ-dysregulated genes; Figure 5G) and were reversed in their expression pattern. Most genes that were upregulated in the EoE transcriptome were downregulated by FICZ, and conversely, many downregulated EoE transcriptome genes were upregulated by FICZ (98 genes of the 156 overlapping genes; Figure 5G). The overlapped 156 genes were enriched for cornified envelope proteins (Figure 5H), suggesting that loss of epithelial differentiation, as observed in EoE biopsies, can be partially reversed by AHR activation.

### IL-13 and IL-4 inhibit OVOL1 activation

We aimed to determine the effect of IL-4 and IL-13, Th2 cytokines with established roles in atopic diseases, including EoE (14, 31, 32), on OVOL1. In unstimulated cells that overexpress OVOL1, OVOL1 was primarily localized to cytoplasmic vesicles that bordered the membranal protein desmoglein 1 (DSG1) (Figure 6A). FICZ stimulation induced nuclear mobilization of OVOL1, whereas IL-4 or IL-13 stimulation prevented the FICZ-induced OVOL1 mobilization to the nucleus in OVOL1-overexpressing cells (Figure 6A). IL-4 and IL-13 also significantly decreased *SPINK7* promoter activity in OVOL1-overexpressing cells that were stimulated with FICZ (Figure 6B-C). Because cells that were differentiated in ALI culture express high endogenous levels of OVOL1, we analyzed the effect of IL-4 and IL-13 on differentiated cells. In unstimulated differentiated cells, OVOL1 was mostly nuclear and remained nuclear after FICZ stimulation (Figure 6D). IL-4 or IL-13 stimulation promoted OVOL1 translocation from the nucleus to the cytoplasm (Figure 3F and Supplementary Movie 1-3). FICZ stimulation significantly increased endogenous *SPINK7* expression (Figure 6E), and IL-13 significantly decreased *SPINK7* expression when the cells were stimulated with FICZ (Figure 6E). As a control for FICZ and IL-13 stimulation, *CYP1A1* and *CCL26* expression were also examined (Figure 6F,G). Interestingly, FICZ stimulation was able to decrease the IL-13–dependent CCL26 expression by 2.7 fold (p < 0.0001; Figure 6G). Transcriptomic analysis of FICZ-, IL-13–, and IL-13 + FICZ–treated cells compared with untreated cells revealed that the 4 groups of treatment were significantly different (Figure 6H and Supplementary Information). The majority of the IL-13–dysregulated genes (55%) overlapped with the IL-13 + FICZ transcriptome (Figure 6I, Supplementary Table 3), whereas less than 10% of the IL-13 + FICZ transcriptome overlapped with the IL-13 transcriptome (Figure 6I, Supplementary Table 3). Addition of FICZ attenuated the IL-13-induced genes, including for key genes such as *CAPN14* (encoding for the cysteine protease calpain 14) and *CCL26* (encoding for the eosinophil chemoattractant eotaxin-3) (Figure 6J). These results suggest that AHR activation counters genes that are modified by IL-13, indicating that treatment with AHR ligands might ameliorate EoE.

**Figure 6.**
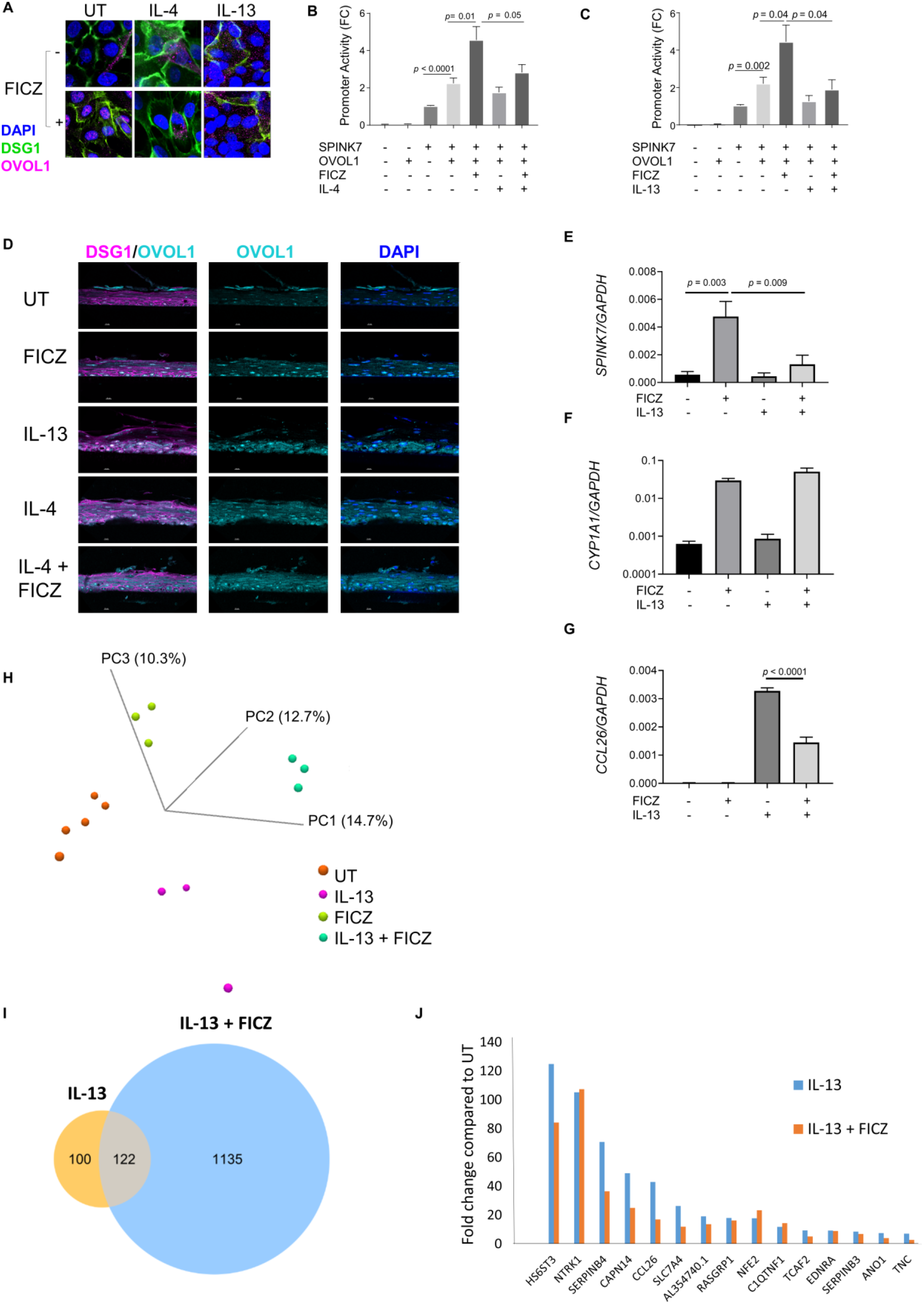
IL-13 and IL-4 prevent OVOL1-dependent *SPINK7* expression. **A**. Representative images of co-immunofluorescence of desmoglein1 (DSG1, green), OVOL1 (pink), and DAPI (blue) stain in OVOL1-overexpressing cells that were either left untreated (UT) or treated over night with IL-4 or IL-13 (100 ng/mL) with or without FICZ (1 µm). Promoter activity in lysates triple-transfected with firefly vector and either *SPINK7*-nLUC or nLUC and either OVOL1 or a control plasmid. Cells were either left untreated or treated with 1 µm FICZ, with or without IL-4 **(B)** or IL-13 **(C). D.** Representative images of co-immunofluorescence of DSG1 (pink), OVOL1 (cyan), and DAPI (blue) stain in cells that were differentiated in the ALI model. Cells were either left untreated or treated with FICZ (1 µm), IL-4 or IL-13 (100 ng/mL), or IL-4 (100 ng/mL) with FICZ (1 µm). mRNA expression of *SPINK7* **(E)**, *CYP1A1* **(F)**, or *CCL26* **(G)** normalized to *GADPH* in cells that were either left untreated or stimulated with IL-13 (100 ng/mL) and/or FICZ (1 µm). **H.** Principal component analysis using a permanova weighted test of dysregulated genes by RNAseq data from cells that were either left untreated or stimulated with IL-13 (100 ng/mL) and/or FICZ (1 µm). **I.** Overlap between the IL-13 transcriptome (IL-13 compared to UT) and IL-13 and FICZ transcriptome (IL-13 + FICZ compared to UT). **J.** The fold change of IL-13 treatment compared to UT and the fold change of IL-13 + FICZ treatment compared to UT of top upregulated genes in the IL-13 transcriptome.

### IL-13–induced *CAPN14* expression depletes OVOL1 protein expression

Because IL-13 is a major driver of epithelial transcriptional changes, we asked if IL-13 can regulate OVOL1 expression. IL-13 stimulation in differentiated esophageal epithelial cells decreased OVOL1 protein expression by 10 fold (p = 0.02; Figure 7A). As a positive control, DSG1 protein expression was decreased by approximately 20% (p = 0.01; Figure 7A). In contrast, *OVOL1* mRNA expression was unchanged between IL-13–stimulated and control cells (Figure 7B). Consistent with the protein level, *DSG1* mRNA decreased in IL-13–stimulated compared to untreated cells (p = 0.018; Figure 7B). These data demonstrate that IL-13 stimulated cytoplasmic retention of OVOL1 and loss of OVOL1 (Figure 4C-D), whereas a nuclear shift of OVOL1 may enable protein stability.

**Figure 7.**
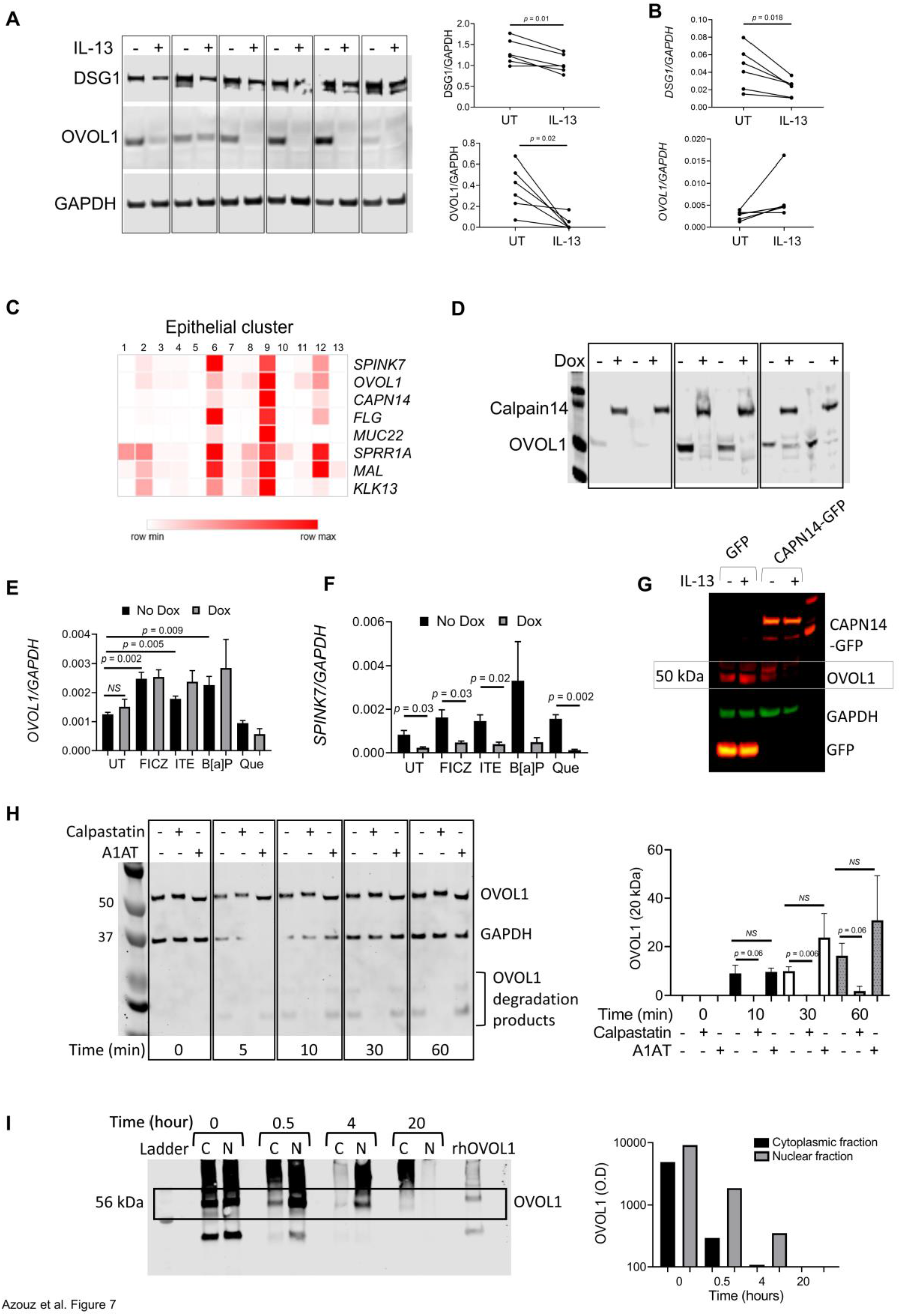
OVOL1 undergoes IL-13–dependent post-translational modifications. **A.** Western blot analysis of OVOL1, DSG1, and GAPDH expression in differentiated EPC2 cells that were either left untreated (UT) or stimulated with IL-13 (100 ng/mL) for 48 h. The graphs on the right show quantification of OVOL1 and DSG1 relative to GAPDH with or without IL-13 treatment from paired UT/IL-13 samples from 6 independent experiments. **B.** qPCR analysis of *OVOL1* and *DSG1* mRNA expression in differentiated EPC2 cells that were either left untreated or stimulated with IL-13 (100 ng/mL) for 48 h; expression is normalized to *GAPDH*. **C.** Heatmap depicting the relative expression of the indicated genes in epithelial clusters on the basis of single-cell RNA sequencing data of dispersed cells from esophageal control biopsies. **D.** Western blot analysis of OVOL1 and calpain 14 expression in differentiated EPC2 cells with inducible expression of *CAPN14* expression. *CAPN14* is fused to a flag tag and is induced by doxycycline (Dox) treatment. qPCR analysis of *OVOL1* **(E)** and SPINK7 **(F)** in differentiated EPC2 with inducible expression of *CAPN14* expression of *CAPN14*; expression is normalized to *GAPDH*. **G.** Western blot analysis of OVOL1 in GFP-overexpressing or CAPN14-GFP– overexpressing cells with or without IL-13 treatment. GAPDH was used as a loading control. Anti-GFP was used for detection of GFP and CAPN14-GFP. **H.** Western blot analysis of protein lysates that were incubated for the indicated times with recombinant OVOL1 protein (100 ng) with or without calpastatin or A1AT. The graph on the right side represents the mean values ± SEM of the measured fluorescent signal of the degradation products of OVOL1, calculated as percent from the measured fluorescent signal of the full length of OVOL1 at time 0, from 3 independent experiments. **I.** Western blot analysis of recombinant human OVOL1-GST (60 ng) that was either left untreated or incubated with cytoplasmic protein fractions (C) or nuclear protein fractions (N) for the indicated times. The graph on the right is a quantification of OVOL1 band intensity (O.D). NS, not significant; UT, untreated.ht is a quantification of OVOL1 band intensity (O.D).

The EoE transcriptome is enriched for proteases and has an imbalance between proteases and protease inhibitors, favoring a proteolytic state (17, 33, 34). Single-cell RNA sequencing analysis that was previously performed by us (35) revealed that *CAPN14* and *OVOL1* are co-expressed in the same epithelial cluster in the esophagus (Figure 7C). The epithelial clusters that co-expressed *OVOL1* and *CAPN14* were enriched for *SPINK7* and corresponded to cells that express differentiation markers (i.e., *FLG*, *MUC22*) and esophageal enriched genes (i.e., *MUC22*, *MAL*, *KLK13;* Figure 7C). The expression of *CAPN14* and *OVOL1* was highly correlated across cells (r = 0.92; Spearman correlation). These findings prompted us to hypothesize that calpain 14 may be involved in OVOL1 post-transcriptional regulation. Notably, CAPN14 is an esophagus-specific protease, encoded by the *CAPN14* gene, which is located in a strongly EoE-associated risk locus (i.e., 2p23) (25, 36). CAPN14 is transcriptionally induced by IL-13 and has been shown to regulate epithelial barrier homeostasis and repair (16, 25). Inducible *CAPN14* expression in differentiated esophageal epithelial cells revealed a marked reduction in OVOL1 protein (Figure 7D). In contrast to OVOL1 protein, *OVOL1* mRNA expression was not affected by inducible *CAPN14* expression (Figure 7E)*. OVOL1* mRNA was induced > 2 fold by AHR ligands even in cells with induced *CAPN14* expression (p = 0.009-0.002; Figure 7E). Induction of *CAPN14* expression decreased *SPINK7* expression by 3.5 fold (p = 0.03; Figure 7F). The expression of *SPINK7* was decreased by 3.5-12 fold after *CAPN14* induction even after administration of AHR ligands (p = 0.03-0.002; Figure 7F). In addition, constitutive expression of CAPN14-GFP decreased the expression of OVOL1 compared to that of control GFP vector transduction (Figure 7G), indicating that the reduction in OVOL1 protein expression resulted from *CAPN14* expression and not as a result of doxycycline treatment. Consistently, IL-13 treatment decreased OVOL1 expression in the CAPN14-GFP–overexpressing cells (Figure 7G). We then tested the hypothesis that a calpain inhibitor would inhibit OVOL1 degradation. Indeed, the calpain inhibitor calpastatin inhibited the degradation of OVOL1 when added to the cell lysates with OVOL1 (Figure 7H). As a control, the serine protease inhibitor alpha-1 antitrypsin (A1AT) did not inhibit the degradation of OVOL1 under the same conditions (Figure 7H).

We hypothesized that nuclear mobilization of OVOL1 by AHR protects OVOL1, whereas IL-13–mediated cytoplasmic retention of OVOL1 promotes degradation of OVOL1 by cytoplasmic proteases, such as calpain 14. Accordingly, we incubated recombinant OVOL1 protein with cytosolic or nuclear extracts; OVOL1 protein degraded more quickly with incubation with cytoplasmic proteins compared to nuclear proteins (Figure 7I). These results indicate that OVOL1 is likely degraded outside of the nucleus and remains relatively stable in the nucleus.

## Discussion

Herein, we identified that delivery of AHR ligands promotes the translocation of OVOL1 to the nucleus, which then induces a transcriptional program of differentiation and barrier function genes, including *SPINK7*. We demonstrate that IL-4 and IL-13 repress this pathway by inhibiting OVOL1 nuclear localization. The cysteine protease calpain 14 (which is induced by IL-13 stimulation (16)), inhibits *SPINK7* expression by enhancing the posttranslational degradation of OVOL1. Therefore, IL-13 can potentially inhibit *SPINK7* expression by 2 mechanisms; first, by inhibiting OVOL1 nuclear translocation, which prevents OVOL1 from binding to its target genes (e.g., SPINK7 promoter) and second, by inducing calpain-14 expression, which in turn degrades OVOL1 and prevents *SPINK7* expression.

Herein, we revealed that the transcription factor OVOL1 controls *SPINK7* expression. OVOL1 is an enriched esophageal transcription factor that is induced during esophageal epithelial differentiation (27, 37). OVOL1 regulates expression of barrier genes such as *FLG* and *LOR* in the skin (28). In addition, variants in the *OVOL1* gene associate with atopic dermatitis, a type 2 allergic disease with strong similarities to EoE, including impaired barrier function (38–40). Our data demonstrates that *OVOL1* depletion decreases *SPINK7* expression and promotes impaired esophageal barrier function and TSLP production. OVOL1 protein expression was lost in esophageal biopsies from patients with EoE compared to controls. Therefore, our findings reveal an altered molecular pathway that is relevant in the disease state. We suggest that OVOL1 has a key role in regulating esophageal homeostasis and immune homeostasis. The potential role of P53 and DNA damage in regulating SPINK7, has been reported (41), calling attention to examining this in type 2 immunity.

We further demonstrated that OVOL1 activity is regulated by AHR. AHR, a member of the basic helix-loop-helix per-Arnt-sim (bHLH/PAS) protein family, reacts to a variety of environmental ligands such as vehicle exhaust and cigarette smoke, dietary compounds (i.e. flavonoids, indole-3-carbinol derivatives, extracts from fruits and vegetables especially cruciferous), products from commensal bacteria, tryptophan metabolism, and importantly, proton pumps inhibitors (42, 43). AHR is trapped in a cytosolic multiprotein complex consisting of heat shock protein 90, tyrosine kinase c-src, and other co-chaperones; translocates from the cytoplasm into the nucleus upon ligand binding; and dimerizes with AHR nuclear translocator (ARNT). The AHR /AHR ligand/ARNT complex recognizes promoters containing xenobiotic responsive elements (XRE), and then activates the transcription of target genes such as phase I and phase II detoxification enzymes (e.g., cytochrome P450 (CYP1A1)) (42). Our data suggest that the AHR pathway is dysregulated in the esophagus of EoE patients compared to control patients, in contrast to AHR levels which were increased in the esophagus of EoE patients compared to control patients, phase II enzymes were markedly decreased (Supplementary Figure 5). These data suggests that alterations in the AHR pathway occurs in EoE patients and can have downstream implications such as alterations in OVOL1 and the OVOL1-regulated transcriptional program.

Interestingly, we provided evidence that Omeprazole, a proton pump inhibitor used for esophageal eosinophilia, can induce *SPINK7* expression by promoting AHR-dependent nuclear translocation and activation of OVOL1. Recently, it has been shown that in 10-50% of patients with EoE, proton pump inhibitors monotherapy can effectively reverse the histologic and clinical features of the disease (44, 45). It has been suggested that the positive effects of proton pump inhibitors on EoE stem from blockade of acid and anti-inflammatory effects, such as inhibition of eotaxin-3 and STAT6 (46). Rochman *et al.* reported that proton pump inhibitor promoted diverse epithelial changes through AHR (47). The findings that the proton pump inhibitor Omeprazole promotes *SPINK7* expression makes it tempting to speculate that part of the positive effects of Omeprazole in EoE may be mediated by *SPINK7* expression and subsequently improved barrier function and esophageal restoration of homeostasis. Our data also demonstrated that another AHR ligand, FICZ, partially inhibits IL-13–dependent eotaxin-3 (*CCL26*) expression.

Allergic diseases are influenced by diverse factors such as diet, infections, exposure to antibiotics and chemicals, microbiome composition and stress, collectively term exposome (the sum of external factors that an individual is exposed to throughout their lifetime) (3, 4, 6, 8, 9, 48). Genetic and epigenetic elements have the potential to influence disease onset (10-12, 49-53). The esophagus directly interacts with and adapts to the external environment. Yet, esophageal exposomal sensors are mostly undefined. AHR is activated in response to multiple environmental ligands and is capable of initiating distinct signaling pathways in response to these signals (29, 30). In this way, AHR senses host/microbiome dysbiosis, dietary compounds, drugs and environmental toxicants and initiates an appropriate cellular response. The data presented here, identify a regulatory network that controls *SPINK7* expression in esophageal epithelial cells in response to environmental signals through AHR and OVOL1. The increase in the prevalence of allergic diseases in the last decades is likely attributed to changes in environmental factors such as crowded living conditions, pollution, hygiene and excessive use of detergents, use of antibiotics and microbiome dysbiosis(3, 4, 6, 48). In addition, an unhealthy western diet and the consequences of the global climate crisis such as poor quality of air, water and food, are further contributing to the increase of allergic diseases. We suggest that these changes in environmental factors are shifting the exposure of healthy AHR ligands (such as degradable natural ligands that are found in fruits and vegetables or bacteria metabolites) and pathogenic AHR ligands that induce negative signals (such as non-degradable exogenous ligands found in pollution). As a consequence of this shift, AHR signaling pathways are altered and subsequently induce susceptibility and sensitization of vulnerable individuals. Notably, Hidaka et al showed that chronic topical application of an exogenous ligand, 7,12-dimethylbenz[a]anthracene, induced atopic dermatitis-like phenotypes in mice but that topical application of the endogenous ligand, FICZ did not (30). Increased prevalence of allergic diseases such as EoE may be a ramification of modified ecosystems impeding AHR-mediated responses.

In conclusion, we have interrogated components of the innate immune system of the esophageal epithelium with an initial focus on the mechanisms that regulate a key checkpoint pro-inflammatory inhibitor SPINK7. We identified elevated histone 3 acetylation marks during cellular differentiation at the *SPINK7* promoter region. We identified binding sites for the C2H2 zinc finger transcription factor OVOL1 in the SPINK7 promoter region and co-localized them with the histone 3 acetylation marks. Furthermore, we found that binding of OVOL1 to the *SPINK7* promoter was required for its activity. Mechanistically, a variety of AHR ligands (e.g., proton-pump inhibitors, dietary compounds, metabolites produced by bacteria and particulate matter), were shown to induce OVOL1 nuclear localization and subsequent *SPINK7* expression. This was particularly notable as PPIs are considered therapeutic for some EoE patients, providing a mechanistic explanation as well as early proof-of-principle for the potential of AHR ligands to be therapeutic for EoE. Furthermore, we demonstrated that the type 2 cytokines IL-4 and IL-13 repressed OVOL1 activation. Additionally, the product of the chief EoE susceptibility locus (2p23) calpain-14 (25), an intracellular regulatory protease induced by IL-13 in esophageal epithelial cells (16), was demonstrated to be involved in post-transcriptional modification of OVOL1 levels. Translational studies identified a marked loss of OVOL1 protein expression in esophageal biopsies of EoE patients compared to control patients. We suggest that AHR serves as an esophageal sensor for environmental signals and has the potential to rapidly control esophageal epithelium fate by altering SPINK7 levels via OVOL1 activation. It has not escaped our attention that our findings have implications for other type 2 allergic diseases such atopic dermatitis, a disease genetically linked to *OVO1* variants, as well as Netherton’s Syndrome, caused by mutations in *SPINK5* (20, 22, 38–40, 54). Given these collective observations, we propose that the homeostasis in esophageal allergic inflammation may be restored by treatment with select AHR ligands. These findings have immediate implication as an AHR ligand (tapinarof) is now in clinical usage for psoriasis and shows promising results in clinical trials for atopic dermatitis (55, 56), and humanized anti-KLK5 antibodies are in development (57). A deeper understanding of the reported findings and their in vivo relevance are warranted.

## Materials and Methods

### Identification of the *SPINK7* promoter

The 4.5-kb region was chosen on the basis of bioinformatics analysis of transcriptional and epigenetic data from ENCODE (Encyclopedia of DNA Elements) and BioWardrobe (Cincinnati Children’s Hospital Medical Center [CCHMC] Epigenetic Database). Cross analysis of these databases have shown that the 4.5-kb region consists of highly conserved sites enriched with histone acetylation marks (H3K27ac) and overlapped with DNase clusters in multiple relevant cellular contexts. The BioWardrobe database (internal unpublished data) has shown that a region of 1.8 kb is enriched with H3K27ac marks at 2 kb upstream of the transcription start site (TSS). The 4.5-kb, non-coding putative promoter sequence (without untranslated region) was obtained from the ENCODE UCSC Genome Browser of the Human genome 2013 database (hg38_dna range), and the coordinates are chromosome 5:148307922-148312422.

### Construction of plasmids

Promoter constructs were created by cloning the immediate 4.5-kb region adjacent to the 5’ TSS of *SPINK7* into the promoterless Nano-luciferase reporter vector pNL1.1-NL (Promega). The 4.5-kb sequence and subsequent constructs were created by using primers with the restriction enzyme sites *KpnI-HF* and *XhoI*. We utilized SnapGene software that employed In-Fusion cloning techniques. Cloning was performed with In-Fusion HD methods (Clonteck, Takara Bio Company). pNL1.1-NL is defined as the empty vector (EV). Post-cloning with the sequence of interest is termed as SPINK7 (4.5 kb). The full-length SPINK7 consists of 4.5 kb, and shorter lengths were defined as SPINK7 1 kb–3 kb from TSS. Mutations of OVOL1 binding sites were performed using QuikChange Lightning site-directed mutagenesis kits (Agilent).

### Generation of CRISPR/Cas9 knockout EPC2 cells

Guide RNA (gRNAs) complementary to the *OVOL1* open reading frame sequences and located directly 5’ of a protospacer adjacent motif (PAM) were identified (*OVOL1* gRNA: 5′-TCTCGCCGCGCTCCTCGTCG-3’ [http://tools.genome-engineering.org (58)]), and the following oligonucleotides (*OVOL1* 5’-CACCGCTCGCCGCGCTCCTCGTCG-3’ and 5’-AAACCGACGAGGAGCGCGGCGAGC-3’) were annealed and ligated into the *BbsI* restriction site of plasmid pX459M2 (obtained from CCHMC Transgenic Mouse and Gene Editing Core Facility) to produce pX459M2-OVOL1g3 and pX459M2-CAPN14g3, respectively. EPC2 cells were transfected with pX459M2, pX459M2-OVOL1g3, or pX459M2-CAPN14g3 using Viromer (Origene) according to the manufacturer’s protocol. Transfected cells were selected; cloned; and gDNA was isolated, amplified, and sequenced as previously described (17). OVOL1 protein expression were determined by rabbit anti-human OVOL1 antibody (Sigma Aldrich).

### Nuclear and cytoplasmic extraction and Western blotting

Proteins from cell cultures were extracted with RIPA buffer (Pierce) with protease and phosphatase inhibitors. Loading buffer (Life Technologies) was added, and samples were heated to 95°C for 5 min, subjected to electrophoresis on 12% NuPAGE Bis-Tris gels (Life Technologies), transferred to nitrocellulose membranes (Life Technologies), and visualized using the Odyssey CLx system (LI-COR Biosciences) with IRDye 800RD goat anti-rabbit (LI-COR Biosciences), and IRDye 680RD goat anti-mouse (LI-COR Biosciences) secondary antibodies. The primary antibodies were rabbit anti-OVOL1 (Sigma Aldrich) or rabbit anti-OVOL1 (LifeSpan Biosciences), mouse anti-HSP90 (Cell Signaling Technology Inc), mouse anti–desmoglein-1 (Sigma Aldrich), and rabbit anti-LaminB1 (Abcam). Blots were quantified using the Image Studio software (LI-COR Biosciences).

To prepare nuclear and cytoplasmic extracts, cells were harvested in cold hypotonic lysis buffer (20 mM Tris-HCl, pH 7.4; 10 mM NaCl; 3 mM MgCl_2_), and the suspensions were incubated on ice for 15 minutes. Following addition of NP-40 to a final concentration of 0.5%, cell suspensions were homogenized by vortexing for 10 seconds and centrifuged at 4°C for 10 minutes at 3,000 RPM to pellet nuclei. Cytoplasmic fractions contained in the supernatant were collected, and nuclear pellets were washed twice with PBS prior to resuspension in extraction buffer (10 mM Tris, pH 7.4; 2 mM Na_3_VO_4_, 100 mM NaCl, 1% Triton X-100, 1 mM EDTA, 10% glycerol, 1 mM EGTA, 0.1% SDS, 1 mM NaF, 0.5% deoxycholate, 20 mM Na_4_P_2_O_7_, purchased from Fisher #FNN0011). Nuclear suspensions were incubated on ice for 30 minutes with vortex every 10 minutes and then centrifuged at 4°C for 30 minutes at 14,000 x *g*. Nuclear proteins contained in the supernatant were collected. Nuclear and cytoplasmic extracts were aliquoted and stored at -80°C.

### Electrophoretic mobility shift assays

Single-stranded oligonucleotides containing the binding sequences of interest were obtained from Integrated DNA Technologies and incubated with either 5’-IRDye 700 labeled or unlabeled complementary strands in annealing buffer (composition) for 5 minutes at 95°C and then allowed to slowly return to room temperature to generate fluorescent and non-fluorescent double-stranded oligonucleotide probes. Fluorescent probe (40 ng) was incubated in binding buffer (Tris, NaCl, poly(dI:dC), NP-40, glycerol) with 250 ng total protein from cell extracts or 120 ng recombinant GST-tagged OVOL1 protein (Abnova #H00005017-P01) and non-fluorescent competitor containing the wild-type or mutant binding site for 30 minutes at room temperature and cross-linked at 120 mJ/cm^2^. Antibody against OVOL1 was added after cross-linking and incubated at room temperature for 15 minutes before loading samples. Binding reactions were resolved on 6% native PAGE gel electrophoresis, and fluorescent probes were detected using a LICOR Odyssey CLx system.

### Statistical analysis

Raw luciferase data were measured as relative luminescent units (RLU), defined by the ratio of Nano-luciferase reporter activity (NL) to the Firefly (FF) activity (NL/FF). Normalized data are defined by the ratio of raw data of the promoter activity (NL/FF) to the average of the EV activity (NL/FF). Statistical analysis was completed with GraphPad PRISM. One-way and two-way ANOVA and t-tests were performed.

## Author contributions

NPA: Conceptually led the study, supervised the study, designed and performed experiments, analyzed data, and wrote the manuscript. AMK: Designed and performed experiments and analyzed data. MR, MP, JMC, and MB: Designed and performed experiments. ATD, DM, AL, and CF: Assisted in experimental procedures. LCK and MTW: Assisted in conceptual design of the study. MER: Conceptually lead the study, supervised the study and wrote the manuscript.

## Acknowledgments

We thank A. Rustgi (University of Pennsylvania) for the human telomerase reverse transcriptase–immortalized EPC2 cell line. We also thank S. Hottinger (CCHMC) for editorial assistance, all of the participating families and the Cincinnati Center for Eosinophilic Disorders, and members of the Division of Allergy and Immunology. Funding: This work was supported in part by NIH R37 AI045898, U19 AI070235; the Campaign Urging Research for Eosinophilic Disease (CURED); the Sunshine Charitable Foundation and its supporters; and Denise and David Bunning.

## Supplementary Information

### Supplementary Materials and Methods

#### Transient transfection and luciferase activity assay

Human esophageal epithelial progenitor cells (EPC2) are immortalized cell lines that were donations from the Dr. Anil Rustgi Lab (Columbia University, NI). EPC2 cells were cultured in Keratinocyte Serum-Free Medium (KSFM). The KSFM medium was supplemented with epidermal growth factor (EGF, 1 ng/mL), bovine pituitary extract (BPE 50 mg/mL), and 1X penicillin/streptomycin (Invitrogen). EPC2 cells were grown for 3-4 days until they reached 80-90% confluency. Cells were then harvested by addition of trypsin/EDTA (Invitrogen) and incubated for 3-5 min at 37^°^C. Next, soybean trypsin inhibitor (250 mg/L in 1X DPBS) was added, and cells were pelleted at 300 x *g* for 5 min. EPC2 cells were seeded on day 0 at 200,000 per 4-well plate in two conditions: low calcium (KSFM medium alone at CaCl_2_ 0.09 mM) and high calcium (KSFM at CaCl_2_ 1.8 mM). Twenty-four hours later (day 1) cells were transiently transfected at >95% density with Opti-MEM (ThermoFisher) and Mirus TransIT-2020 (Mirusbio, Madison, WI) according to the manufacturer instructions. We used 2 µL of TransIT-2020 per 500 ng of construct DNA and 50 ng Firefly DNA (1:20 dilution). Cells were co-transfected with 1000 ng pNL1.1 containing the *SPINK7* promoter sequence or promoterless empty vector (EV) pNL1.1-NL with 50 ng Firefly luciferase pGL3 vector (Promega). Firefly, a PGL3 vector (Promega), was employed as an internal control for transfection. On day 2, cells were incubated at 37°C, and the medium was changed with the appropriate calcium conditions. On day 3 (48 h post-transfection), Nano and Firefly luciferase activities were measured in relative light units (RLU) by luciferase assay NanoDLR kit (Promega) via Synergy 2 and Synergy H1 Multi-Mode Microplate Reader (BioTek Instruments, Winooski, VT). In order to control for transfection efficiency, Nano-luciferase activity was normalized to Firefly-luciferase, and then the activity was normalized to control promoterless transfected cells for each sample transfection to account for variance per well. All assays were conducted in triplicate in 3 independent experiments.

#### Endogenous *SPINK7* expression

To determine endogenous *SPINK7* expression, EPC2 cells were plated at high density (250,000 cells/well in a 48-well plate) in KSFM media with 1.8 mM CaCl_2_. For plating at low density, 250,000 cells/well were grown in a 6-well plate in KSFM media with 1.8 mM CaCl_2_. RNA was isolated with Quick-RNA Micro-prep (Zymo; Irvine, CA). ProtoScript First Strand cDNA Synthesis kit (NEB; Ipswich, MA) was employed according to the manufacturer instructions to obtain RT-PCR data. For air-liquid interface (ALI) differentiation, cells were grown as previously described (17, 59). Briefly, 150,000 cells/well were plated in a transwell system with 24-well plate. After 48 h, media was replaced with a high calcium media (1.8 mM CaCl_2_). On day 8, the media was aspirated from the top chambers; on day 12, the cells were stimulated; and on day 14, the cells were harvested.

#### mRNA extraction and quantitative RT-PCR

Total RNA was isolated from cells with the RNeasy mini kit (Qiagen) according to the manufacturer’s protocol. For RNA sequencing experiments, RNA was treated with On-Column DNase Digestion kit (Qiagen) according to the supplied protocol. cDNA was synthesized with the ProtoScript synthesis kit (New England Biolabs). qPCR was performed using a 7900HT Fast Real-Time PCR system from Applied Biosystems (Life Technologies) with FastStart Universal SYBR Green Master mix (Roche Diagnostics Corporation) by using the following primer sets: *GAPDH* (forward 5′-TGGAAATCCCATCACCATCT, reverse 5’-GTCTTCTGGGTGGCAGTGAT), *SPINK7* (forward 5′-GCATTTACAAGAAGTATCCAGTGGT, reverse 5’-TCGTGAAGAAACTGAACTCTTCC), *DSG1* (forward 5’-TGGCTACATTTGCAGGACAA, reverse 5’-CGGTTCATCTGCGTCAGTAG), *OVOL1* (forward 5’-CAAAAGAGGTCCCCAGAACA, reverse 5’-AGGAGCCTTCCTCTCAGGTC), *CYP1A1* (forward 5’-GACAGATCCCATCTGCCCTA, reverse 5’-GGTTGATCTGCCACTGGTTT), *CCL26* (forward 5’-CTCCCAGCGGGCTGTGATATTC, reverse 5’-GGGTCCAAGCGTCCTCGGAT).

#### 3’ RNA sequencing and data processing

RNA sequencing was performed with high-quality RNA (RNA integrity number > 8) by using the QuantSeq 3’ mRNA-Seq Library Prep Kit FWD for Illumina (Lexogen, Vienna, Austria, catalog no 015.96). Libraries were subjected to quality control and concentration measurements at the Gene Expression Core at Cincinnati Children’s Hospital Medical Center (CCHMC). Libraries were diluted to final concentrations of 5 nM and sequenced on a HiSeq 4000 Illumina sequencing machine at the Genomics and Cell Characterization Core Facility at the University of Oregon with 100- to 150-bp–length single reads. Data analysis and visualization were performed using the CLC Genomics Workbench, version 12.0.3 (Qiagen, Hilden, Germany) as described previously (47). Differentially expressed genes were defined by fold change and statistical filtering and clustered, as indicated in the figure legends. Gene ontology (GO) enrichment analysis, which uses statistical methods to determine functional pathways and cellular processes associated with a given set of genes, was performed with the ToppGene suite (60) (https://toppgene.cchmc.org/). Unless otherwise indicated, differentially expressed genes were used as the input for GO analysis. Venny (https://bioinfogp.cnb.csic.es/tools/venny/) was used to intersect gene lists. As indicated, expression data were intersected with the EoE transcriptome, and the list of 1,607 significantly dysregulated transcripts was identified by comparing gene expression in the biopsy specimens of 10 patients with active EoE (9 of which were irresponsive to the proton pump inhibitor [PPI] treatment) with that in normal controls by the RNA sequencing analysis (61).

#### Analysis of epithelial clusters

Expression data of selected genes according to epithelial cluster was obtained from next-generation single-cell RNA sequencing of human esophageal biopsies deposited under GSE201153 (62). Clustering of epithelial cells was described previously (35).

#### *OVOL1* gene silencing by shRNA

Lentiviral shRNA vectors against OVOL1 (MISSION shRNA, Sigma-Aldrich, clone NM_004561, TRCN00000257410, TRCN0000229665, TRCN0000229664, and TRCN0000229666) and a control vector that targets no known mammalian genes (SHC002 SIGMA MISSION® pLKO.1-puro Non-Mammalian shRNA Control) were used. EPC2 cells grown in KSFM media were transduced. Twenty-four hours after transduction, cells were selected for stable integration using puromycin (1 μg/mL). After ALI differentiation, gene silencing efficiency of target vectors in transduced cells was assessed by quantitative PCR relative to that of cells transduced with non-silencing control (NSC) shRNA.

#### Immunofluorescence

Cells that were grown in ALI cultures on top of a membrane were fixed with 10% formalin and embed in paraffin to preserve the architecture. Formalin-fixed, paraffin-embedded (FFPE) samples were sectioned and de-paraffinized using xylene and then subjected to graded ethanol washes. Heat-induced epitope retrieval in sodium citrate buffer (10 mM sodium citrate, 0.05% Tween 20, pH 6.0) was used. Slides were blocked in 1X phosphate-buffered saline (PBS) with 10% goat serum for 1 h followed by a 1-h incubation at room temperature in the following primary antibodies: rabbit anti-human OVOL1 (Sigma Aldrich) and mouse anti–human DSG1 (Santa Cruz Biotechnology). Slides were then washed, incubated for 1 h at room temperature in secondary antibodies (donkey anti– rabbit Alexa Fluor 488, donkey anti-Rabbit Alexa Fluor 568, and goat anti–mouse Alexa Fluor 594 [Thermo Scientific]) and mounted with 4’,6-diamidino-2-phenylindole (DAPI) Fluormount-G (Novus Biological). Images were obtained using the NIKON A1RSi confocal microscope.

### Supplementary Figures

**Supplementary Figure 1. Regulation of the *SPINK7* promoter. A.** Promoter activity in lysates co-transfected with promoter less nLUC and firefly vector that were grown in the indicated concentrations of CaCl2. Promoter activity was determined by nLUC measurements relative to firefly measurements and normalized according to the promoter less nLUC measurements. Promoter activity is presented as relative luminescence units (RLU). **B.** Promoter activity in lysates of cells that were grown in 0.09 mM of CaCl_2_ and co-transfected with nLUC constructs that contain either 0, 1, 2, 3, 4 or 4.5 kb of the *SPINK7* promoter sequence and firefly vector. Promoter activity was determined by nLuc measurements relative to firefly measurements and normalized according to the promoter less nLUC measurements. Data are mean ± SEM.

**Supplementary Figure 2. Expression of OVOL1. A.** Western blot analysis of OVOL1 in control or OVOL1-overexpressing EPC2 cells. β-actin was used as a loading control. **B.** Expression of *OVOL1* mRNA in non-silencing control cells (control) and *OVOL1*-silenced cells (*OVOL1* KD). Data are mean ± SEM.

**Supplementary Figure 3. Generation of *OVOL1* knockout EPC2 cells. A.** A chromatogram depicting the genomic DNA sequence of EPC2 cells in the vicinity of the sequence targeted for CRISPR/Cas9-mediated editing. The box indicates the location of the PAM sequence. **B.** Prediction of the protein sequences of *OVOL1* knockout cells and control cells according to their genomic sequence. Black text indicates amino acids that match the wild type (WT) protein sequence. Blue text indicates amino acids that deviate from the WT protein sequence.

**Supplementary Figure 4. OVOL1 expression in EoE and control esophageal biopsies.** Western blot analysis of OVOL1 expression in control and eosinophilic esophagitis (EoE) biopsies. The graph on the right shows the OVOL1 expression relative to HSP90.

**Supplementary Figure 5. Dysregulation of phase II enzymes in EoE.** Expression of *AHR*, *NQO1, HMOX1*, and *HMOX2* in biopsies from 10 patients with eosinophilic esophagitis (EoE) compared with 6 control patients (control). Data are mean ± SEM.

**Supplementary Movie 1. Nuclear expression of OVOL1 after cellular differentiation.** Reconstituted, three-dimensional confocal images of representative co-immunofluorescence staining of DSG1 (green), OVOL1 (pink), and DAPI (blue) staining in cells that were differentiated in air-liquid interface (ALI) culture.

**Supplementary Movie 2. IL-13 inhibits OVOL1 nuclear localization after differentiation.** Reconstituted, three-dimensional confocal images of representative co-immunofluorescence staining of DSG1 (green), OVOL1 (pink), and DAPI (blue) staining in cells that were differentiated in air-liquid interface (ALI) culture and were treated with IL-13 (100 ng/mL).

**Supplementary Movie 3. IL-4 inhibits OVOL1 nuclear localization after differentiation.** Reconstituted, three-dimensional confocal images of representative co-immunofluorescence staining of DSG1 (green), OVOL1 (pink), and DAPI (blue) staining in cells that were differentiated in air-liquid interface (ALI) culture and were treated with IL-4 (100 ng/mL).

**Supplementary Table 1. List of candidate transcription factors (TFs) that may regulate SPINK7 expression.** Bioinformatics analyses of TFs that are predicted to bind the *SPINK7* promoter sequence, TFs that are dysregulated in eosinophilic esophagitis (EoE) compared to control, TFs that are induced during epithelial differentiation, and TFs that are enriched in the esophagus (37).

**Supplementary Table 2. AHR transcriptome.** Differentially regulated genes following FICZ (1 µM) 18-h treatment compared to untreated (UT) cells, as indicated by fold change of > │2│. Data obtained on the basis of RNA sequencing data from EPC2 cells that were either left untreated or treated with FICZ (1 µM).

**Supplementary Table 3. FICZ, IL-13, and FICZ + IL-13 transcriptomes.** Expression (TPM values) of genes in control cells (untreated cells), FICZ (1 µM) treatment, IL-13 (10 ng/mL) treatment, and IL-13 (100 ng/mL) + FICZ (1 µM) treatment.

